# Rats Sniff off Toxic Air

**DOI:** 10.1101/739003

**Authors:** Haoxuan Chen, Xinyue Li, Maosheng Yao

## Abstract

Breathing air is a fundamental human need, yet its safety, when challenged by various harmful or lethal substances, is often not properly guarded. For example, air toxicity is currently monitored only for single or limited number of known toxicants, thus failing to fully warn against possible hazardous air. Here, we discovered that within minutes living rats emitted distinctive profiles of volatile organic compounds (VOCs) via breath when exposed to various airborne toxicants such as endotoxin, O_3_, ricin, and CO_2_. Compared to background indoor air, when exposed to ricin or endotoxin aerosols breath-borne VOC levels, especially that of carbon disulfide, were shown to decrease; while their elevated levels were observed for O_3_ and CO_2_ exposures. A clear contrast in breath-borne VOCs profiles of rats between different toxicant exposures was observed with a statistical significance. Differences in MicroRNA regulations such as miR-33, miR-146a and miR-155 from rats’ blood samples revealed different mechanisms used by the rats in combating different air toxicant challenges. Similar to dogs, rats were found here to be able to sniff against toxic air by releasing a specific breath-borne VOC profile. The discovered science opens a new arena for online monitoring air toxicity and health effects of pollutants.

**TOC:** 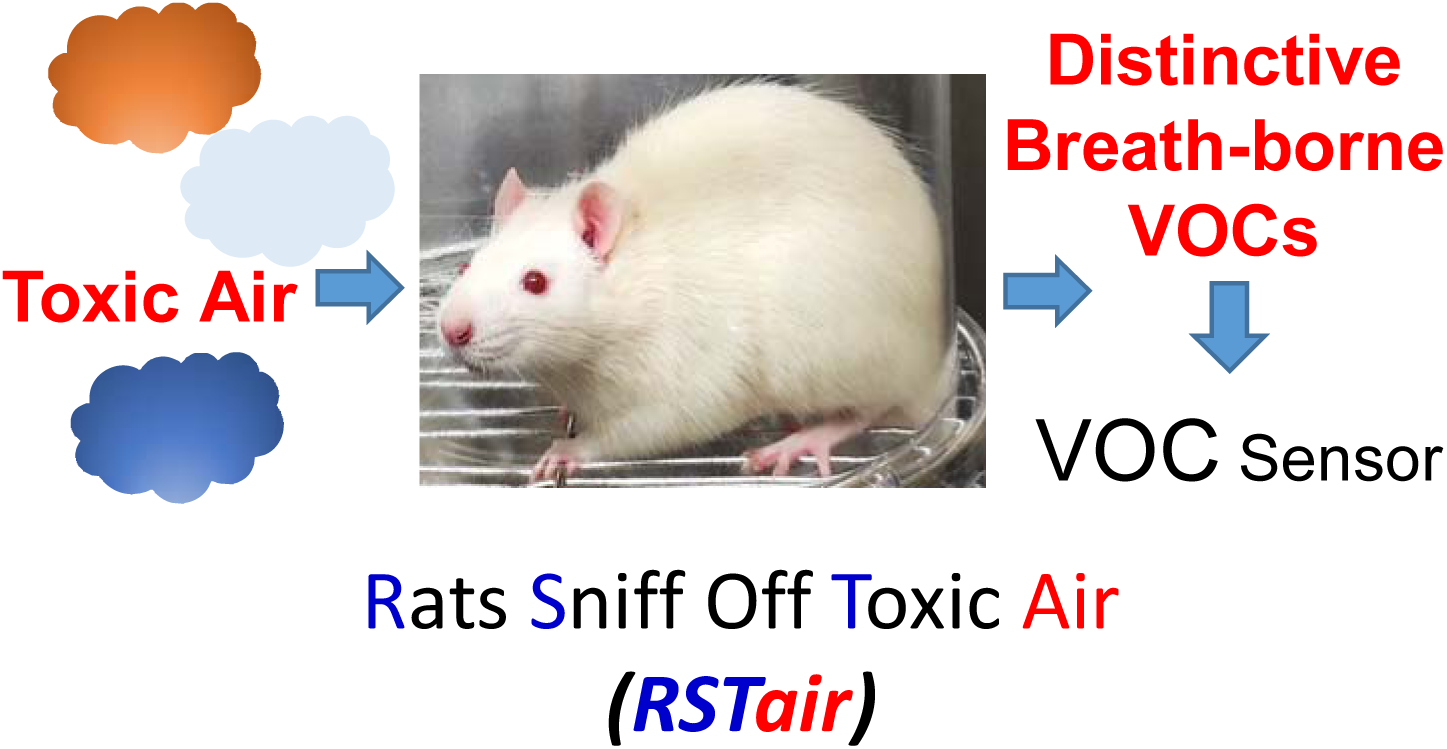

## 1. Introduction

Polluted air is a complex mixture of various pollutants, including particulate matter (PM), biologicals, and also gaseous substances such as O_3_ and NOx. Inhaling these pollutants can cause a variety of serious health problems: respiratory and cardiovascular diseases, and even death. ^1-2^ For example, PM alone was shown to have resulted in 4.2 million deaths in 2015. ^2-3^ Exhibiting a positive correlation with daily mean mortality, ground ozone exposure resulted in decreased lung function and airway inflammation. ^4-5^In the meantime, exposure to pathogenic bioaerosols including bacteria, fungi, virus, et al in the air can cause respiratory infections with grave human and economic costs.^6-9^ Apart from these, there is a growing risk of terrorist attacks by intentionally releasing biological and chemical agents into the air to cause substantial civilian casualties. ^10-12^ Apparently, inhaling unsafe air has become an increasing health concern. Yet, in many high profile events, in addition to the public sectors, the air being inhaled is not readily protected or properly guarded. Real-time monitoring of air toxicity is of great importance, which however is a long-standing significant challenge in the field.

For monitoring hazardous substances in the air, a variety of real-time online monitoring methods have been previously developed or tested for individual pollutants such as the PM and other chemicals. ^13-16^While for bioaerosols, the adenosine triphosphate (ATP) bioluminescence technology, surface-enhanced Raman spectroscopy (SERS), bioaerosol mass spectrometry (BAMS), ultraviolet aerodynamic particle sizer (UVAPS) as well as silicon nanowire (SiNW) biosensor were investigated and attempted over the years. ^17-19^ It is well known that these existing or developed technologies are mostly restricted to either single agent or overall microbial concentration levels without identifying species. In addition, airborne pollutants and toxicity could vary greatly from one location to another ^20-21^, thus presenting location-specific air toxicity and health effects. Current epidemiological or toxicological methods involving data analysis or animal and cells experiments cannot provide in situ air toxicity information, accordingly failing to represent the response at the time of exposure since biomarker levels evolve over time. ^22^ In addition, under certain scenarios (high profile meetings or locations) a rapid response to air toxicity needs to be in place in order to protect the interests. However, the response time is very demanding for an immediate effective counter-measure, for example, usually several minutes can be tolerated. ^10, 23^ In many air environments, multiple hazardous pollutants could also co-exist even with unknown ones at a particular time, which makes protecting the air rather difficult, if not impossible, using current technologies of species level detection and warning.

Previously, olfactory receptors of mouse cells for odors ^24^, immune B cells ^25^for pathogen detections, and silicon nanowire sensor arrays for explosives^26^ were studied. Recently, a breath-borne biomarker detection system (dLABer) integrating rat’s breath sampling, microfluidics and a silicon nanowire field effect transistor (SiNW FET) device has been developed for real-time tracking biological molecules in the breath of rats exposed to particulate matter (PM).^27^ The dLABer system was shown to be able to online detect interleukins-6 (IL-6) level in rat’s breath, and capable of differentiating between different PM toxicities from different cities using the biomarker level. However, as observed in the study the production of protein biomarkers could significantly lag behind the pollutant exposure, thus falling short of providing a timely warning against toxic air. Nonetheless, exhaled breath is increasingly being used in biomarker analysis in both medical and environmental health studies. ^28-30^ In addition to protein biomarkers, a large number of volatile compounds (e.g., nitric oxide, carbon monoxide, hydrocarbons) have been also studied to assess health status and even developed for clinical diagnosis.^31-32^ For example, ethane and n-pentane detected in the breath were linked to the *in vivo* level of lipid peroxidation and oxidative stress ^33^; breath-borne acetone was shown to correlate with the metabolic state of diabetic patients.^34^ In addition, exhaled VOCs have been used for the diagnosis of asthma, lung cancer and other diseases.^35-37^ Undoubtedly, breath-borne VOC has emerged as a promising biomarker for health or environmental exposure monitoring.

Inspired by the dog sniff for explosive, the work here was conducted to investigate if we can use breath-borne VOCs from living rats to real-time monitor toxic air. Particularly, we wanted to investigate: 1) When rats are exposed to air toxicants, whether the VOC species and concentration in the exhaled breath change? If yes, how long does the change need to occur? 2) Are there any specific exhaled VOCs in response to different toxicants exposure including both chemical and biological agents? 3) To develop an online air toxicity analyzing system via real-time monitoring of exhaled VOCs of rats. The work here has demonstrated a great promise for online air toxicity monitoring using the method developed, and opened a new arena for studying health effects of air pollution.

## 2. Materials and methods

### 2.1 Rat breeding

The Jugular Vascular Catheterizations (JVC) rat model described in our previous study was used in this work.^38^ Weighing 200–240 g at an age of 10 weeks, a total of 20 male Wistar rats with jugular vein catheterization operation and a 100 mL/min inhalation rate were purchased from a local provider (Beijing Vital River Laboratory Animal Technology Co., Ltd.). With about 1 centimeter out of the skin, a flexible sterile catheter was embedded into the jugular vein and fixed on the back with staples. Under a 12 h light/12 h dark cycle, all the rats were raised in an animal care facility naturally with a normal chow diet. After 1 week of acclimation, the rats were randomly divided into 5 groups (4 rats in each group) for the exposure of different air toxicants. All animal experiments were approved by the Institutional Review Board of Peking University and relevant experiments were performed in accordance with the ethical standards (approval # LA2019294).

### 2.2 Preparation of toxic air

In this work, four different exposure toxicants (ricin, endotoxin, ozone and carbon dioxide) and indoor air (as a background control) (5 groups) for rats were prepared for the exposure experiments. Ricin was extracted from the seeds of castor produced in Xinjiang, China, and prepared by Institute of Microbiology and Epidemiology, Academy of Military Medical Sciences in Beijing. The endotoxin was purchased from Associates of Cape Cod, Inc., USA. The ricin and endotoxin suspensions were prepared by vigorously vortexing 40 μg of ricin or 50 ng endotoxin per ml deionized (DI) water for 20 min at a vortex rate of 3200 rpm (Vortex Genie-2, Scientific Industries Co., Ltd., USA). Detailed information about ricin preparation and exposure can be also found in another work.^39^ Here, ozone was generated by an ozone generator (Guangzhou Environmental Protection Electric Co., Ltd., China) using corona discharge method. The ozone was further diluted with indoor air for rat exposure experiments, and the ozone concentration in the exposed chamber was approximately 5 ppm. Carbon dioxide was purchased from Beijing Haike Yuanchang Utility Gas Co., Ltd., and diluted 20 times with indoor air to a concentration of about 5% (5×10^4^ ppm). The major objective of this work was to study whether there will be specific VOC release by rats when challenged with different toxicants, not specific dose-response for each toxicant. Nonetheless, the selection of specific exposure levels for different toxicants was provided and discussed in Supporting Information.

### 2.3 Rats sniff off toxic air (RST_air_) system and experimental procedure

To investigate whether we can use breath-borne VOCs from living systems to real-time monitor toxic air, we have developed the system named as *RSTair* (Rats Sniff Off Toxic Air). As shown in Figure 1 and Figure S1 (Supporting Information), the system is composed of four major parts: toxicant generator, exposure chamber, exhaled breath sampling and online VOC analysis. Indoor air was used as carrier gas for generating toxicant aerosols (ricin and endotoxin) using a Collison Nebulizer (BGI, Inc., USA) or diluting toxicants gas (ozone and carbon dioxide). The toxicant aerosol or toxicant gas was introduced together with indoor air into the exposure chamber at a flow rate of 1 L/min. As shown in Figure S1, a cage was used as the exposure chamber which can allow rat’s feces and urine to fall from below quickly so as to reduce their influences on VOC analysis. In addition, teflon tubes and vales were also applied to reducing adsorption loss of VOCs. For ricin and endotoxin, they are small molecules and relevant particle loss on the aerosolization tubing could be negligible. When performing the experiments, the rats were first placed in the exposure chamber. Indoor air was continuously introduced into the chamber at a flow rate of 1 L/min for 10 minutes, then followed by each of tested toxicants together with indoor air at the same flow rate for about 10 minutes to conduct exposure tests.

**Figure 1.**
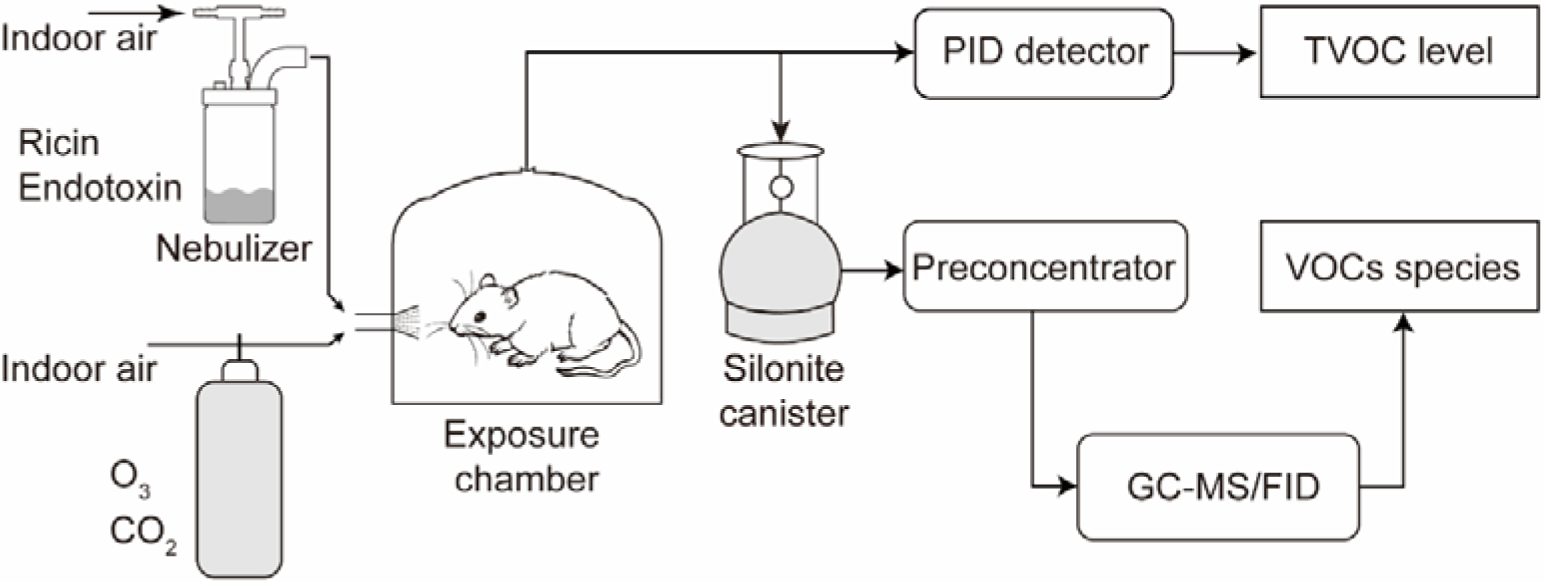
Rats sniff off toxic air (*RST*air) system: different toxicants were introduced or aerosolized into the chamber for exposure. The VOCs from the rat placed in the chamber before and after the toxicant exposure for 10 min were analyzed by the PID directly and also by GC–MS/FID method coupled with a VOC sampler silonite canister. During the VOC collection and measurements by the PID, the toxicant exposure was terminated. Each time only one rat was placed into the chamber. For each toxicant, the experiment was independently repeated four times with a different rat each time.

For each toxicant experiment, the TVOC was first measured for the control (rat with indoor air at a flow rate of 1 L/min), then followed by the toxicant aerosol/gas exposure (rat + toxic air) at the same flow rate for 10 min. Here, the photoionization detector (PID) (MOCON, Inc., USA) coupled with an air pump was used to real-time monitor TVOC at a flow rate of 0.6 L/min. After the exposure, the TVOC was continuously measured again using the PID sensor, reaching a relatively stable level approximately within 7-8 min. In the meantime, VOC samples were also collected both before (control) and after the toxicant exposure. A 3.2 L Silonite canister (Entech Instruments, Inc., USA) was used to collect VOC samples at the flow rate of 0.8 L/min at least 10 min after the exposure test, and the VOCs species were further analyzed using a gas chromatograph-mass spectrometry/flame ionization detection (GC–MS/FID) system (Agilent Technologies, Inc., USA). A total of 117 VOC species were screened and quantified for all exposure tests using the GC-MS system. During the VOC sample collection, the PID sensor was switched to measure the indoor air TVOC levels. The detailed descriptions of the working principles of PID and VOC species analysis by GC–MS/FID are provided in Supporting Information. In this work, both TVOC and VOC species in the exposure chamber were analyzed for all the experiments: 1) when the rats were not in the chamber (background air); 2) rats in the chamber (before toxicant exposure) and rats in the chamber (after toxicant exposure). The air flows in and out of the exposure chamber were the same both for TVOC measurements and VOC collection. Based on the dimensions of the exposure chamber, the residence time of pollutants in the chamber was about 5 min under the experimental conditions, e.g., the air flow rate was about 1 L/min. We collected VOC samples at least 10 min after the toxicant exposure, therefore remaining secondary pollutants produced, if any, during the exposure tests such as O_3_ will be flushed out by the indoor air. For each toxicant, the above experiments were repeated four times with a different rat each time. Each rat (total four) either for toxicant or control exposure group experienced the exposure only once, not repeated exposures. For each of four tested toxicants, the same experiments were repeated. All tests were performed inside a Class 2 Type A2 Biological Safety Cabinet (NuAire, Plymouth, MN), and all exposure toxicants were ventilated out via the built-in and lab ventilation system.

### 2.4 Blood microRNA detection

Right before and 20 min after each 10-min toxicant exposure, 0.75 mL blood samples from rats were taken through the catheter using sterile syringes with 23G flat-end needles and kept at −20 °C for microRNA analysis. The blood microRNAs in the blood samples such as miR-125b, miR-155, miR-146a, miR-21, miR-20b, miR-210, miR-122, and miR-33 were analyzed using a RT-qPCR array (Wcgene Biotech, Inc., China). Total RNAs in the blood samples, including microRNAs, were extracted using a Trizol reagent (Sigma Aldrich, Inc., USA) according to the manufacturer’s instructions. Subsequently, the purified RNAs were polyadenylated through a poly(A) polymerase reaction and was then reversed-transcribed into complementary DNA (cDNA). TIANGEN® miRcute Plus miRNA First-Strand cDNA Kit (Code No. KR211) was used in the reverse transcriptional reaction system of total 10□μL, including 5□μL□2x miRNA RT Reaction Buffer, 1□μL miRNA RT Enzyme Mix and 4□μL RNA sample. The reaction conditions are 40 °C for 60 mins and 95 °C for 3 mins. The cDNA was quantified in real-time SYBR Green RT-qPCR reactions with the specific microRNA qPCR Assay Primers. TIANGEN® miRcute Plus miRNA qPCR Kit (SYBR Green) (Code No. FP411) was used in the qPCR reaction system of total 10□μL, including 5□μL□2x miRcute Plus miRNA PreMix (SYBR&ROX), 0.2□μL Forward Primer, 0.2□μL Reverse Primer, 1□μL 50X ROX Reference Dye, 1□μL DNA Sample and 2.6□μL ddH_2_O. The cycling conditions were 95 °C for 15 min, followed by 40 cycles at 94 °C for 20s, 60 °C for 15s and 72°C for 30s. The primers used for qPCR are presented in Table S1 (Supporting Information).

## 3. Statistical analysis

In this study, the TVOC levels for all samples detected by the PID sensor were not normally distributed, while their TVOC change rates were. The TVOC change rate was calculated by dividing the TVOC level after exposure using the TVOC level before exposure. Therefore, the Mann-Whitney rank sum test was used to analyze the differences in TVOC levels before and after each toxicant exposure, and t-test was used to analyze the differences in TVOC change rates between each toxicant exposure group and the control group (indoor air). For individual VOC concentrations by GC-MS/FID, the paired t-test was used to analyze differences for each VOC species before and after the exposure. The software Canoco 4.5 was used to visualize the VOC profile distance and relatedness between the samples of different groups using the principal component analysis (PCA). Besides, the concentrations of micro-RNAs in blood samples from different rat groups were determined by RT-qPCR. For the group exposed to carbon dioxide, blood samples were only taken from two rats (before and after the 10-min exposure) because of the catheter blockage (unable to draw bloods) for the other two. For the other three groups, blood samples were obtained for all four rats. The differences between micro-RNA levels in blood samples before and after the exposure in one group were analyzed using a paired t-test (data exhibited a normal distribution) or Wilcoxon signed rank test (data did not follow a normal distribution). The outliers for normally distributed data were examined and eliminated by a Grubbs test. All the statistical tests were performed via the statistical component of SigmaPlot 12.5 (Systat Software, Inc., USA), and a *p-value* of less than 0.05 indicated a statistically significant difference at a confidence level of 95%.

## 4. Results and discussion

### 4.1 TVOC monitoring for the toxicants exposure

As described in the experimental section, four toxicants (ricin, endotoxin, ozone and carbon dioxide) and indoor air (as a background control) were used for inhalation exposure in rats. For each group, the TVOC levels of only 3 rats (PID instrument/software failure for one out of 4 rats) were shown in Figure 2. The indoor air background TVOC was found to be less than 0.04±0.02 ppm. After one rat was placed in the exposure chamber, the TVOC level in the cage was shown to first gradually increase, then reach a relatively stable level after about 500 seconds. The TVOC level before the exposure (rat + indoor air) when one rat was in the chamber was about 2 ppm, except for the CO_2_ group it was about 0.5-0.8 ppm (These differences, if any, applied to both before and after exposure in one test, thus presenting no influences on the same experiments). The differences in TVOC levels for indoor air exposures (different times: “before” and “after”, but the same indoor air) were small (the average change rate was about −4%±1.4% (95% confidence interval)), although the Mann-Whitney Rank Sum Test showed that for each of the rats, the difference (over some time for the indoor air) was significant (*p-value*<0.001). The background indoor air TVOC levels were 1.22-5.1% (detected) of the chamber TVOC with rats together with indoor air or other toxicants). The fluctuations were taken into account for each toxicant exposure test. Nonetheless, the fluctuations, if any, from background indoor air had minor impacts on the TVOC levels measured for rats’ exposure tests. The change rates (n=3) of the control group (indoor air) then served as the reference for other toxicant exposures in the statistical analysis. During the indoor air experiment, the rats were seen to carry out normal life activities in the exposure chamber, and correspondingly the TVOCs in the chamber were shown to remain relatively stable.

**Figure 2.**
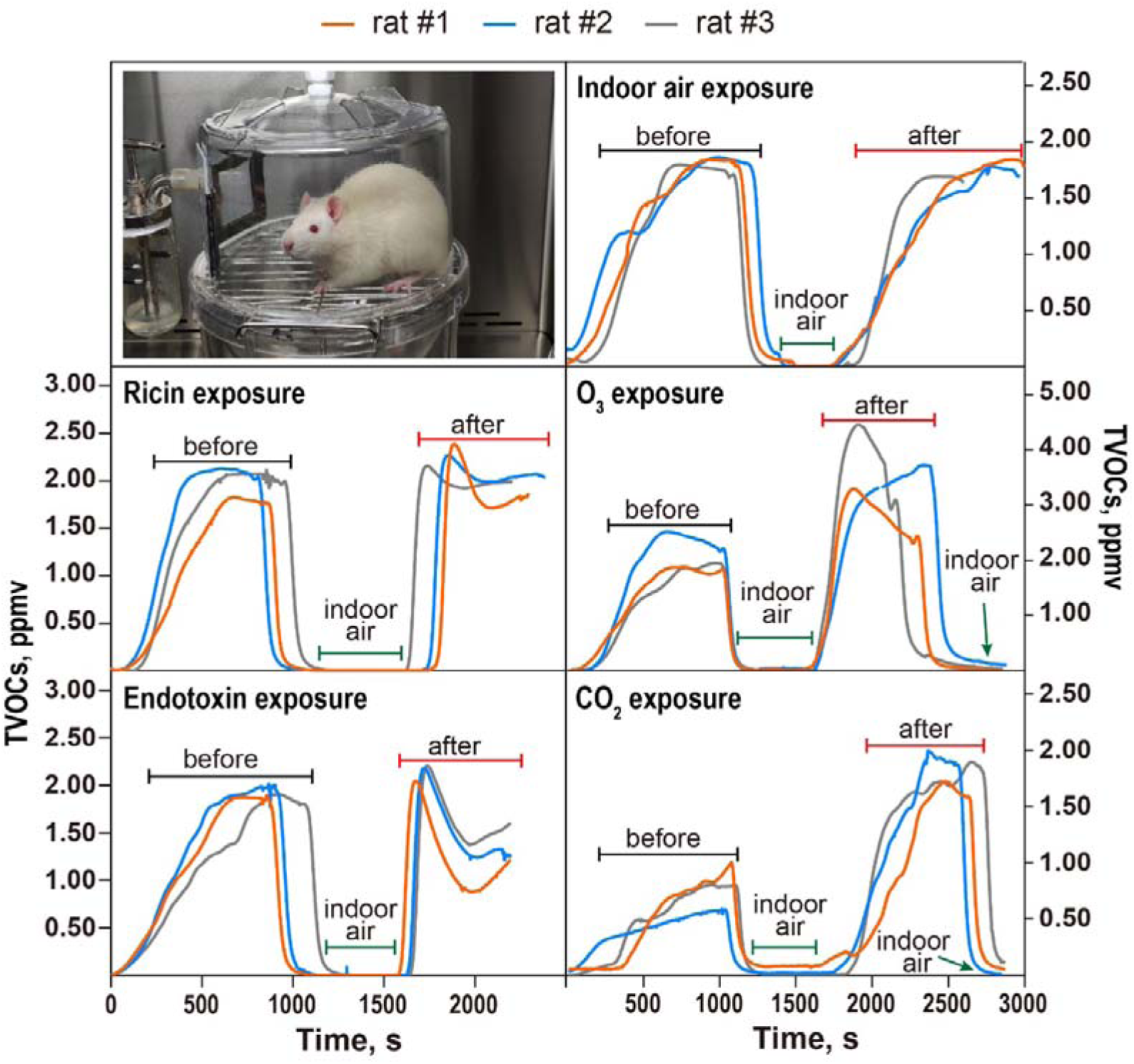
Real-time continuous measurements of exhaled TVOC levels in the chamber when rats were exposed to different toxicants via inhalation for 10 mins: Indoor air, ricin, endotoxin, O_3_ and CO_2_. During the exposure processes, the PID sensor was turned off. Data lines (measurement time was 1000 s) represent results from three individual rats (#1, #2, #3) before or after exposure to each of the air toxicants (aerosolized amounts described in the experimental section) tested. Each exposure test was independently repeated with four rats from the same group (PID instrument/software failure for one rat).

In contrast, the TVOC levels were shown to vary greatly with different toxicant exposures as shown in Figure 2. For example, when rats were exposed to the aerosolized ricin, the TVOC level was observed to exhibit an average change rate of −3%±1.6. Compared to the control group (indoor air + rat before the exposure) shown in Figure 2, the difference of the TVOC change rate was not significant for the ricin exposure (t-test, *p-value*=0.426). For ricin exposure, its concentration (40 μg/mL suspension aerosolized) might be too low in aerosol state after the aerosolization from the liquid to produce a detectable response from the rats. It was previously reported that 0.1 mg/mL ricin solution was used for aerosolization and subsequent exposure to mice (Ref # 5, Supporting Information). After 30 minutes ricin aerosol exposure, the exposed mice became poisoned. The concentration of ricin used in this experiment was lower and the exposure time was shorter, therefore the toxicity reaction may be mild. In contrast to the ricin exposure, however we observed a different phenomenon for the endotoxin (50 ng/mL aerosolized) tests as shown in Figure 2. Upon the endotoxin exposure, the TVOC level was observed to first increase, and then decreased to a level that was about 21-46% below the pre-exposure level after four minutes (Mann-Whitney Rank Sum Test, all *p-values*<0.001). Compared to the control group (indoor air + rat), the difference of the TVOC change rate was statistically significant for endotoxin (t-test, *p-value*=0.0147). The observed differences from the ricin and endotoxin exposures could be due to different mechanisms initiated by different substances involved. Ricin is derived from plant, while endotoxin is from Gram-negative bacterial membrane. They could interact differently with relevant human respiratory or other body cells. Nonetheless, for both ricin and endotoxin, they were probably causing health effects by immuno-toxicity, while O_3_ and CO_2_ both induce harm first by chemical manners.

After exposure to gaseous toxicants such as ozone and carbon dioxide, the levels of TVOCs in the exposure chamber with rats were observed to have increased significantly, as observed in Figure 2. As can be seen from the figure, the TVOC levels has increased for about 44-110% for ozone and about 109-265% for carbon dioxide exposure (Mann-Whitney Rank Sum Test, all *p-values*<0.001). The t-test showed that differences of the TVOC change rates of both ozone and CO_2_ exposures compared to the control group (indoor air) were statistically significant (*p-value*=0.0219 and 0.0296, respectively). These data indicated that rat exposure to both ozone and CO_2_ has resulted in significant elevations of TVOCs, suggesting rats were actively responding to the exposure challenges. The behavior observation from a video also indicated that rats after the exposure to O_3_ seemed to be suffering from the challenge (Video S1, Supporting Information).

### 4.2 Changes in exhaled VOCs after exposure to different toxicants

Using GC-MS/FID method, a total of 31 different VOCs out of 117 screened were detected and shown in Figure S2 (Supporting Information) for background indoor air with and without rats. Among detected VOCs as shown in Figure S2 (Supporting Information), the VOCs with the highest concentrations in indoor air were n-hexane, ethyl acetate and acetone. When rats were placed in the exposure chamber (one rat at each time), the most abundant VOC species was detected to be acetone, which was about 4 times more than that of the indoor air background. Statistical tests found that the concentrations of ethylene and ethane in the chamber containing one rat were significantly lower than those of the background (paired t-test, *p-value*<0.05), which in part could be due to the air dilution by the rat’s breath. Namely, when rats’ breath with specific higher or lower VOC species replaced equivalent indoor air inside the chamber, diluting effects for higher indoor VOC species and enhancing effects for lower indoor VOC species could take place.

Differentiations of VOC species from the rat’s exhaled breath under different toxicants exposures were also shown in Figure 3. There were no significant differences in the concentrations of any VOCs before and after the exposure for the control group, i.e., indoor air (t-test, *p-value*=0.05). This suggests that indoor air is relatively less toxic at a level that is unable to detect a VOC change. In contrast, specific VOC species had experienced significant changes when rats were exposed to ricin, endotoxin, O_3_ and CO_2_ as observed from Figure 3. For example, exposure to ricin caused significant higher concentration of ethyl acetate (183% higher), while lower concentration of carbon disulfide (22% lower). As shown in Figure 3, after the endotoxin exposure process, concentrations of five VOC species: ethane, acetone, cyclopentane, carbon disulfide and methylcyclopentane were shown to be significantly different with those of before the exposure (t-test, all *p-values*<0.05). As can be seen from the results of the ozone exposure group in Figure 3, the concentrations of propionaldehyde, pentane, 2-butanone, hexane and 2-methylpentane exhibited significant differences before and after exposure (t-test, all *p-values*<0.05), in which all the VOCs except 2-methylpentane were elevated. In comparison, rat exposure to CO_2_ resulted in acetone level increase by 34% (t-test, *p-value*=0.0016). These data suggest that exposure to different toxicants had led to production of different profiles of VOC species in addition to their level changes.

**Figure 3.**
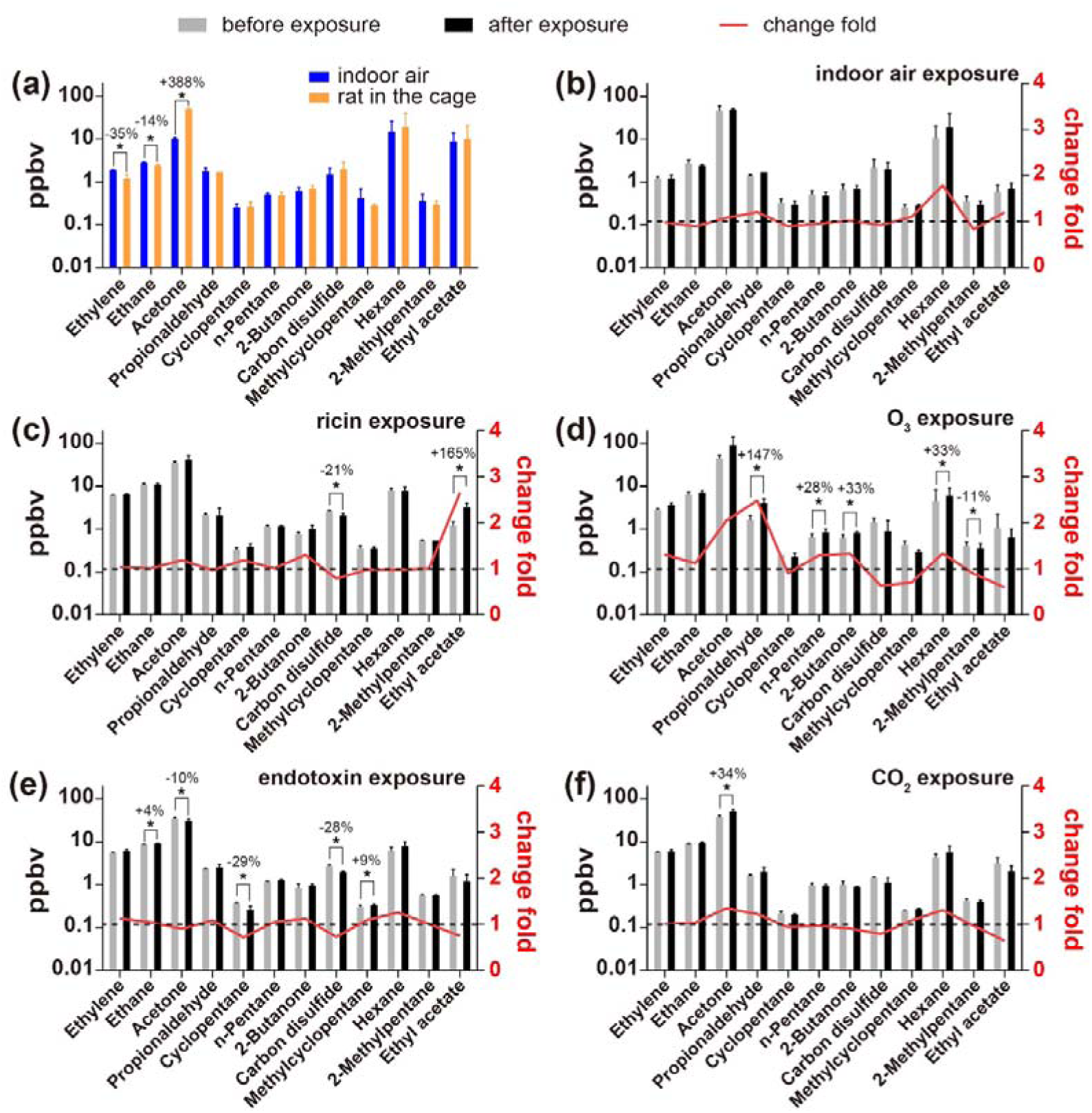
Differentiations of VOC species from indoor air and those from the rats’ exhaled breath under different air toxicity with exposure to ricin, O_3_, endotoxin and CO_2_. The red lines show that the average change ratios of every toxicant calculated by the level after the exposure divided by the level before exposure (right axis). The dotted line is the baseline with a change ratio of 1. Percentages refer to specific VOC percentage change before and after the exposure. Values represent averages and standard deviations from three different rats. “*” indicates a significant difference at *p-value*<0.05.

### 4.3 Detection of micro RNAs in Blood Samples

To further explore the VOC response mechanism of rats to toxicants exposure, microRNAs (miRNA) in the blood samples were examined by an RT-qPCR assay. Fold-changes in microRNA levels after toxicants exposure were shown in Table S2 (Supporting Information). The level of miR-33 in the blood of rats was shown to be significantly lower than that before ricin exposure (*p-value*<0.05); after exposure to ozone, miR-146a level in the blood samples of rats was significantly higher than those before the exposure (*p-value* <0.05), while miR-155 was significantly lower than that before the exposure (*p-value* <0.05). For other microRNAs as listed in Table S2, the changes seemed to be insignificant (t-test, *p-values*>0.05).

As observed from Figure 4, PCA results revealed a clear contrast in breath-borne VOC profiles of rats between different toxicants exposures. The VOC profiles of the ozone exposure group was very different from that of the control group (indoor air) and the VOCs profile of the ricin exposure group was the closest to that of the control group, which agreed with TVOC level and VOC species profiles obtained above. Overall, the experiments showed that rats responded differently to different toxicants by releasing different VOC species owing to different mechanisms of toxicity: ozone caused significant increases in various breath-borne VOCs; while endotoxin exposure generally decreased the releasing of VOCs; and ricin and carbon dioxide exposure resulted in one or two significant VOC species changes. In general, the results of qualitative and quantitative analysis by the GC-MS/FID method agreed with the TVOC level monitored by the PID sensor.

**Figure 4.**
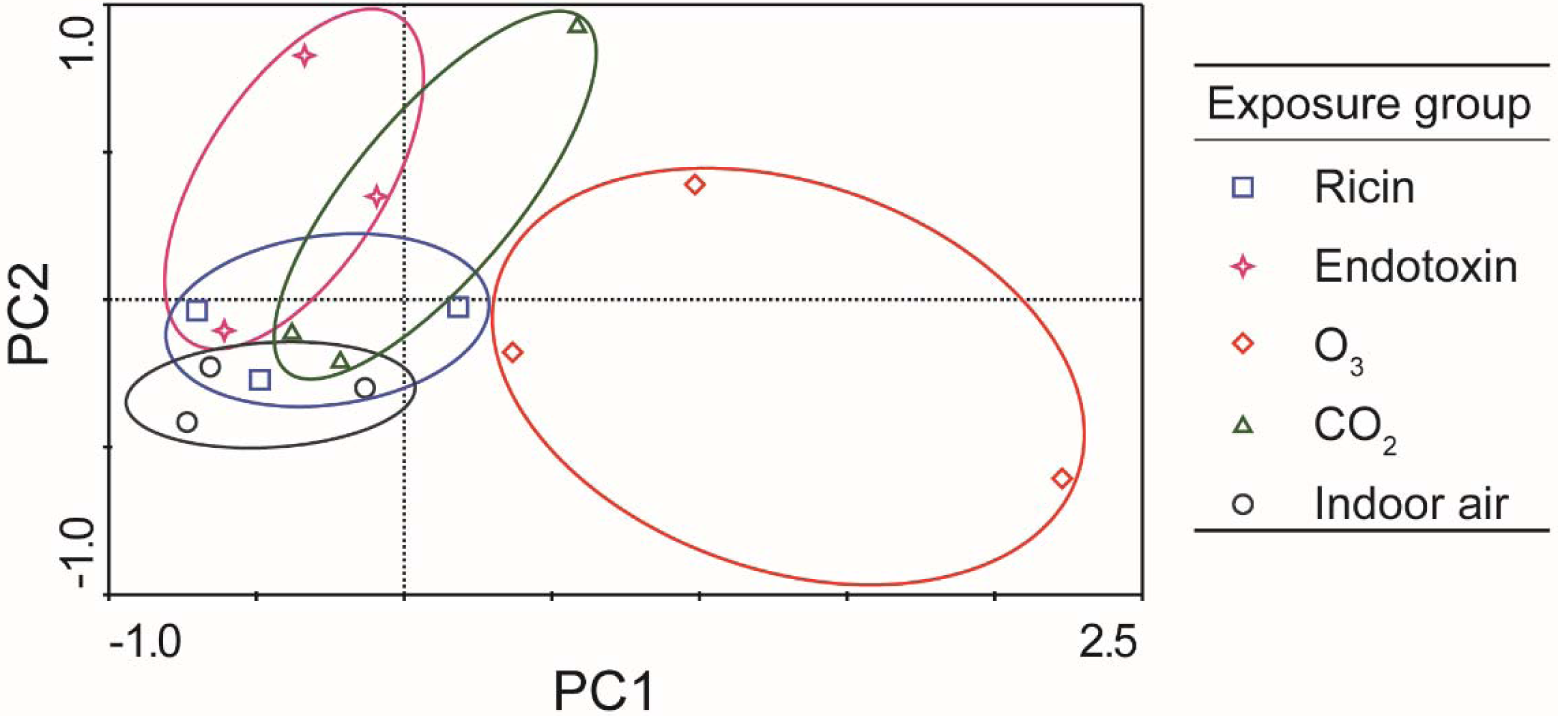
PCA ordinations of exhaled breath-borne VOCs profiles under exposures to different toxicants: ricin, endotoxin, O_3_, CO_2_ and control (indoor air). PC1 and PC2 are the first and second principal components. The VOCs species involved in the PCA analysis were the 12 species which were shown to have undergone changes after each toxicant exposure. Data presented in the figure were from three independent rats exposed to each toxicant.

As observed from Figure 3, exposure to ricin caused 183±143 % higher concentration of ethyl acetate, while 22±8 % lower concentration of carbon disulfide. It was previously reported the concentration of ethyl acetate was significantly higher in exhaled breath from people with cancer compared to the healthy group.^40,41^ In addition, *in vitro* experiments have shown the human umbilical vein endothelial cells (HUVEC) can produce ethyl acetate, which is presumably generated by a reaction of ethanol with acetic acid.^42^ It was demonstrated that ricin is not only responsible for the ricin intoxication through ribosomal inactivation and subsequent inhibition of protein synthesis and cell death, but also shows endothelial toxicity by acting as a natural disintegrin binding to and damaging human endothelial cells.^43^ Therefore, the toxicity of ricin on the endothelial cells might be the source of the higher concentration of ethyl acetate observed in this work. As a disease biomarker, carbon disulfide was observed in the exhaled breath.^44-45^ Recently, it was suggested that the carbon disulfide may be generated endogenously and play a role as a bioregulator. ^46^ Here, we observed that exposure to both ricin and endotoxin resulted in lower levels of breath-borne carbon disulfide compared to the control.

For CO_2_ and endotoxin exposure, the observation for acetone was the opposite as shown in Figure 3. Acetone in exhaled breath was widely investigated in many studies as an important biomarker related to blood glucose and diabetes.^34,47^ Acetone is produced in the fatty acids metabolism by hepatocytes via decarboxylation of excess acetyl coenzyme A (Acetyl–CoA), and then oxidized via the Krebs cycle in peripheral tissue.^48^ As shown in Figure 3, the acetone level increased by 34±9% as a result of CO_2_ exposure, suggesting CO_2_ caused hypoxia in rats, and led to increased respiration from rats. These increases in acetone level corresponded to TVOC level increase as determined by the PID sensor after the exposure to CO_2_. However, when exposed to endotoxin, the acetone level in the exposure chamber decreased by about 10±6%, indicating that the respiration of the rats may be attenuated by the exposure of endotoxin. Clearly, the involved mechanisms by which endotoxin and CO_2_ cause health effects to rats could be very different.

As observed in Figure 3, the increase of ethane level by endotoxin exposure suggested that lipid damage was induced by oxidative stress in the rat’s body since ethane is acknowledged as a marker of lipid peroxidation and described to be generated by peroxidation of ω-3 polyunsaturated fatty acids.^33^ In addition to ethane, the increase of methylcyclopentane level as shown in Figure 3 might also be the result of the endotoxin exposure. Endotoxin has been shown to trigger inflammation through its interaction with the TLR4/CD14/MD2 receptor and then initiates a signal cascade. This reaction correspondingly results in the activation of transcription factor such as NF-κB leading to the production of pro-inflammatory cytokines and type 1 interferons (IFNs), and finally results in systemic inflammatory response syndrome.^49^ In general, in terms with the average fold changes, the concentration of total VOCs in the exhaled breath was relatively lower after the endotoxin exposure, which agreed with the results of TVOC obtained by the PID sensors.

In addition to these biologicals, we have also shown that exposure to chemicals such as ozone and CO_2_ also resulted in in-vivo changes in VOC levels. From the fold changes of various VOC species such as propionaldehyde, N-pentane, 2-Butanone, and Hexane, the ozone exposure has resulted in an overall increase of VOCs in rats’ exhaled breath, which agreed with the TOVC monitoring shown in Figure 2 by the PID sensors. It was previously indicated that increase in propionaldehydes, further products of lipid peroxidation, indicated more severe oxidative damage in rats following exposure to ozone.^50^ Here, we also showed that after exposure to ozone changes in specific breath-borne VOCs and also certain miRNA regulations were detected. Ozone was described as a strong oxidizing agent, and can cause intracellular oxidative stress through ozonide and hydroperoxide formation.^51^ The mechanism of ozone oxidative damage involves the activation of Nrf2, heat shock protein 70, and NF-κB, thus increasing expression of a range of proinflammatory cytokines such as TNFα and interleukin 1β, and chemokines such as interleukin 8.^52^ The results above show that regardless of toxicant types breath-borne VOCs from rats experienced *in vivo* changes.

Results of miRNAs from rats’ blood as shown in Table S4 revealed different mechanisms by rats when exposed to various types of toxicants. miRNAs are short non-coding RNA sequences that regulate gene expression at the posttranscriptional level; and many miRNAs have already been identified to influence physiological processes such as immune reaction, adaptation to stress, and widely investigated in environmental exposure studies.^53^ Among these microRNAs, miR-125b, miR-155, miR-146a, and miR-21 are mainly shown to regulate oxidative stress and inflammatory processes *in vivo*, and widely investigated in air pollution related studies.^53-54^ For example, among them, miR-155 has a positive regulation function, and the other three are negative regulation.^55-56^ However, in this study, the ozone exposure resulted in decreased levels of miR-155 (t-test, *p-value*<0.05), which are different from previous reports in the literature. ^55^ The difference could be due to different exposure toxicant, i.e., ozone used here, leading to acute lung damages compared to mild airway inflammation or asthma problems in other studies. Previous studies have shown that miR-20b and miR-210 are hypoxia regulators in animals, and miR-122 and miR-33 are mainly responsible for regulating lipid metabolism and glucose metabolism in the body.^57-58^ However, these microRNAs, except for the decreased level of miR-33 and increased level of miR-146a caused by ricin and ozone exposure, respectively, were shown to have not undergone significant changes in this study after the exposures to four different toxicants (*p-values*=0.05). The possible reason may be that microRNAs act as post-transcriptional regulators by degrading mRNA or inhibiting their translation, thus failing to respond in a timely manner during short-term exposure (blood samples taken 20 min after the 10-min exposure). To further understand the problem, a yeast model is currently being used to fully investigate the mechanisms underlying the miRNA regulation when exposed to the toxicants used here. Nonetheless, these results here reflected that exposure to the toxicants led to specific miRNAs either up- or down-regulated. On the other hand, the results here also suggest, especially for those no changes observed for the miRNAs, higher toxicant level or specific longer time might be needed to allow miRNA regulation change to occur.

To address the major objective of this work, we repeated a total of 20 times using 5 different exposure agents such as ozone, CO_2_, ricin, endotoxin together with the background indoor air as a control, and 20 different rats. Clearly, above results indicate that when rats are exposed to toxic substances their certain metabolic activities are immediately affected, i.e., these exposures promoted or inhibited specific VOC productions. Based on the results we obtained from this work, the following VOC emission mechanisms of rats when exposed to different toxicants are proposed and illustrated in Figure 5. Previously, it was suggested that VOCs are produced during the normal metabolism in the body; while pathological processes, such as metabolic disorders, can also produce new species of VOCs or alter the levels of existing VOCs.^59^ Therefore, cell or tissue injuries caused by external toxicants exposure also can alter the exhaled VOCs profile by disturbing the normal process.^60^ The exact toxic effect mechanism as observed from this work could vary from one toxicant to another. For some pollutants such as endotoxin and ricin, there are specific receptors to recognize them and then start the chain of responses or reactions.^49,61^ Among these various mechanisms, the ROS (reactive oxygen species) and oxidative stress are recognized to be the central and the common mechanism in various forms of pathophysiology, as well as the health effect of various air pollutants including ambient particulate matter (PM). ^1, 62-63^ Oxidative stress is essentially a compensatory state of the body and can trigger redox-sensitive pathways leading to different biological processes such as inflammation and cell death.^51,64^ For example, the strong oxidants such as ozone might cause oxidative stress through direct effects on lipids and protein^65-66^, which mostly caused the generation and release of hydrocarbons and aldehydes, such as ethane, ethylene, and propionaldehyde.^50,67^ While carbon dioxide tends to make the redox balance tilted toward the reduction side by reducing oxygen supply and thus influencing the energy metabolism in cells.^68-70^ For ricin and endotoxin exposure, the underlying mechanisms seem to be different from ozone and CO_2_, and they could cause oxidative stress indirectly through the activation of intracellular oxidant pathways. Nonetheless, all toxicants share a common effect of disrupting the redox balance, and thus interfere with normal biochemical reactions or cause material damages in cells, accordingly changing the VOC profile released into the breath. As discussed above, in this work, the VOCs profile of rats changed significantly after exposure to different toxicants. Therefore, regardless of toxicant types, breath-borne VOCs from the rats seem to be capable of serving as a proxy for real-time monitoring air toxicity. It was reported about 100 years ago that three mice were also carried on all British submarines for sensing small leakage of gasoline as the rats could squeak to notify the crew (Figure S3, Supporting Information). Some previously detected VOCs such as hexane, pentane, acetaldehyde, butanone, and acetone from rat’s exhaled breath ^71, 72^ were also detected in this work. Nonetheless, due to variations in rat breed, experimental conditions, VOC sampling and analysis methods, it is rather difficult to exactly compare the VOCs from rat’s exhaled breath across different studies. There might be some breath-borne VOC species not identified yet for the rats used here. Interestingly, rat L6 skeletal muscle cells cultivated with α-MEM containing 10% FCS were shown to release 7 VOC species and uptake 16 VOC species.^73^ Similarly, the increased VOCs detected here from rat’s exhaled breath might be released from lung cells or other impacted ones by the exposure. On another front, previous work showed that changes were also detected in breath-borne VOCS from people with upper respiratory infections compared to the health individuals (Ref #7, Supporting Information). All these studies and data support the results and validity of our work from various aspects. For improving *RST*_air_ performance, GC-MS can be also replaced using the Proton Transfer Reaction-Mass Spectrometry (PTR-MS) for fast online VOC species analysis. By using this discovered fundamental science, the invented *RSTair* system here showed its great promise of revolutionizing the air toxicity monitoring, and providing significant technological advances for air security in related fields such as military defense, customs, counter-terrorism and security assurances for important events or special locations.

**Figure 5.**
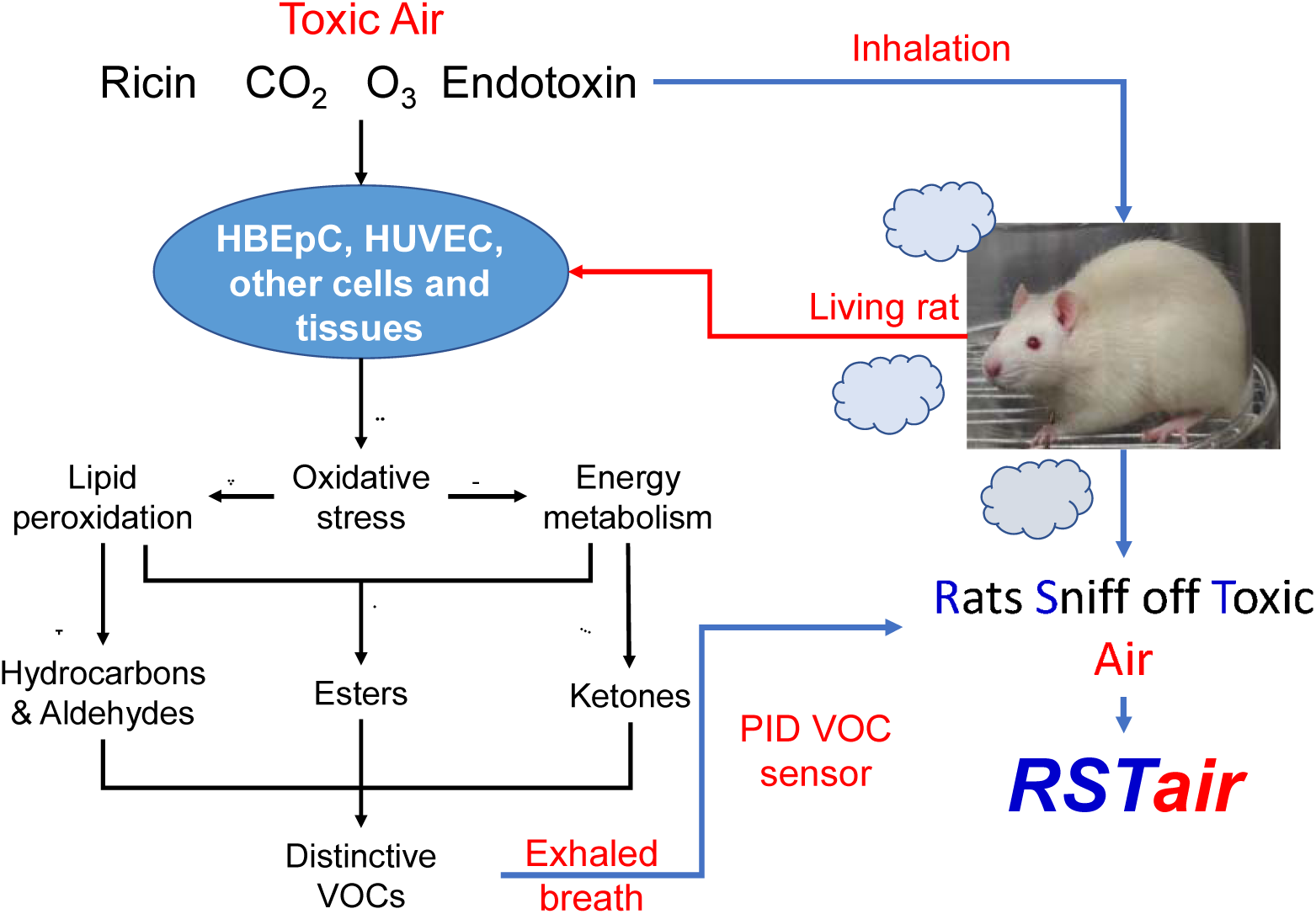
Proposed mechanisms of toxic effects and VOCs releasing in rats when exposed to the environmental toxicants via inhalation. The black arrows represent the toxic effects of different toxicants and the possible pathways of VOCs generation. The blue arrows stand for the principle and working process of the invented *RST*air system for real-time air toxicity monitoring. The corresponding references cited are: ➀ ^59^; ➁ ^66^; ➂ ^63^; ➃ ^48^; ➄ ^42^; ➅ ^48, 70^.

## Supporting information

rat exposed to ozone

## Supporting Information

Measurements of exhaled VOCs by PID and GC-MS/FID, Selection of exposure levels for different toxicant, experimental setup for *RSTair* system, indoor air background VOCs, Photo of UK postcard for three mice carried on British submarines, Primers used for RT-qPCR analysis of microRNA and mi-RNA expression level changes after exposure, video of rat after exposed to ozone were provided as Supporting Information.

## Acknowledgements

This study was supported by the NSFC Distinguished Young Scholars Fund Awarded to M. Yao (21725701), and the Ministry of Science and Technology (2016YFC0207102).

