## Supporting info for "Rats Sniff off Toxic Air"

October 2019

**Supporting Information**

***Measurements of exhaled VOCs by PID and GC-MS/FID***

The working procedure of the PID sensor is to first ionize the VOCs gas with a high-energy UV lamp, and then the ionized fragments are collected by the ion chamber to generate a current signal. The signal in general is proportional to the concentration of the target VOCs. In this study, the PID sensor is assembled with the oil-free pump and pre-filter and a signal encoder, and the signal is transmitted and displayed in real time. The sensor was calibrated using 1 ppm isobutylene prior to use.

Meanwhile, as mentioned above air samples were also collected by a 3.2 L Silonite canister (Entech Instruments, Inc., USA) at the flow rate of 0.8 L/min for the qualitative and quantitative VOC species detection. The air samples were then transferred into the pre-concentrator (7200, Entech Instruments, Inc., USA) and analyzed by gas chromatography (GC) together with Mass Spectrometry (MS) and Flame Ionization Detector (FID) (5975C/7890A, Agilent Technologies, Inc., USA) within 12 hours. The GC-MS/FID analysis work was provided by Tianjin Zhongfei Huazheng Testing Technology Co., Ltd, and is briefly described as follows: the Entech 7200 system was used to pre-concentrate the sample, including enrichment of the sample, and removal of water and carbon dioxide from the sample. Then, an Agilent 5975C/7890A gas chromatography mass spectrometer was used to qualitatively analyze sample according to the standard mass spectrometry library and the mixed gas standard compound. The multi-point calibration working curve was developed by the external and internal standard method for quantitative analysis. The external standard gases used are: mixed standard gas of 57 Photochemical Assessment Monitoring Stations (PAMS) gas, 12 kinds of aldehyde and ketone gases and 47 TO-15&17 gases (including aromatic hydrocarbons, halogenated hydrocarbons and oxygenated organic compounds). The internal standard gases are mixtures of bromochloromethane, 1,4-difluorobenzene, deuterated chlorobenzene, and 1-bromo-3-fluorobenzene. The above standard gases were purchased from Scott Specialty Gases, USA.

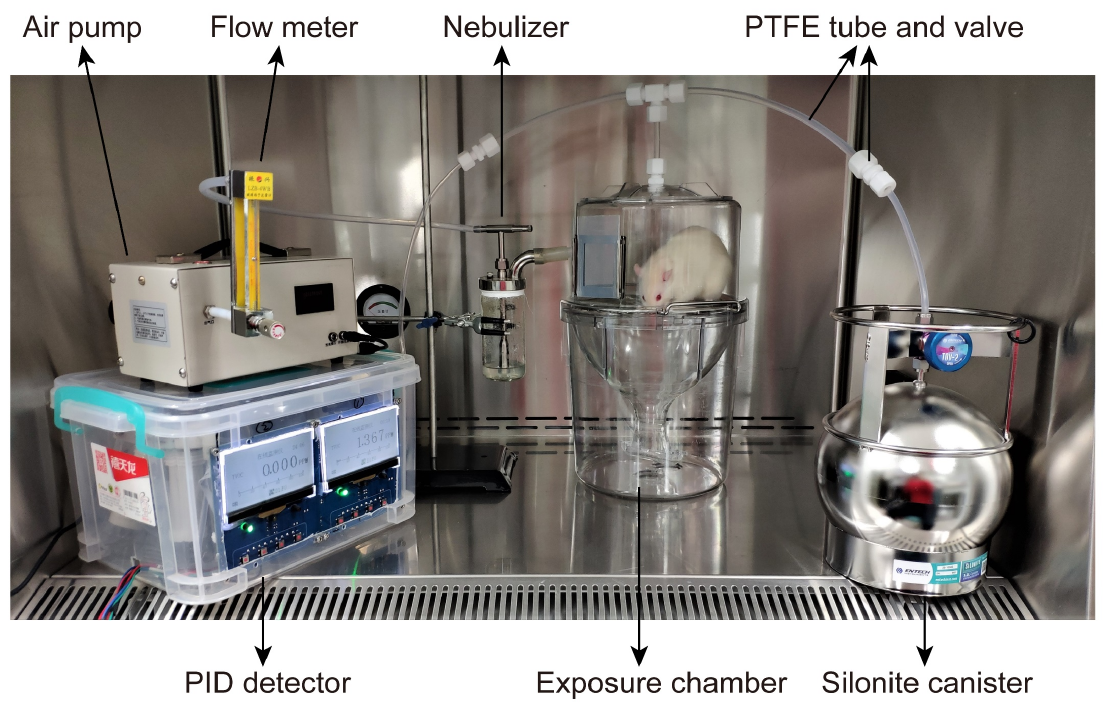

**Figure S1.** Experimental setup (*RST*air) for measuring total VOCs from the rats with and without different toxicant exposures. For each toxicant, at least three rats were tested.

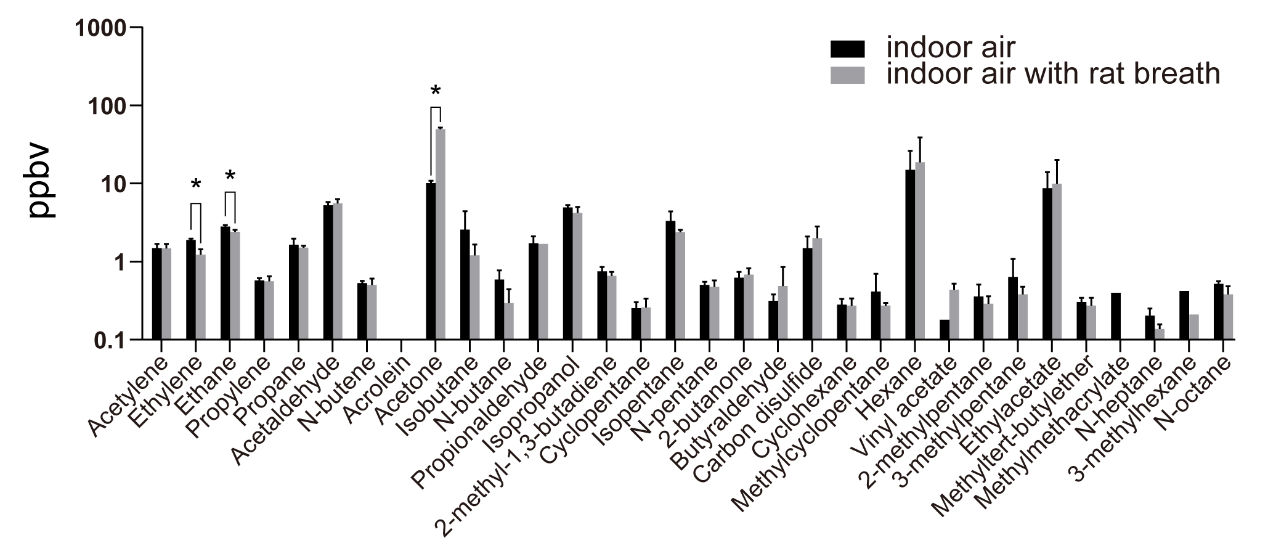

**Figure S2.** Differentiations of VOC species from indoor air and those from rats’ exhaled breath. The data represent average levels of three independent experiments, and error bars stand for the standard deviations. “*” indicates a statistically significant difference at *p-value* <0.05.

**Table S1** Primers used for RT-qPCR analysis of microRNA. The reverse primers are included in the TIANGEN® miRcute Plus miRNA qPCR Kit (SYBR Green) (Code No. FP411).

| microRNA | Forward primers |
| --- | --- |
| MIR-122-5P | GAGTGTGACAATGGTGTTTG |
| MIR-125B-5P | CCTGAGACCCTAACTTGTGA |
| MIR-146A-5P | GAGAACTGAATTCCATGGGTT |
| MIR-155 | TAATGCTAATCGTGATAGGGGTT |
| MIR-20B | AGTGCTCATAGTGCAGGTAG |
| MIR-210 | CCTGCCCACCGCACACTG |
| MIR-21-5P | AGCTTATCAGACTGATGTTGA |
| MIR-33 | GCATTGTAGTTGCATTGCA |

**Table S2** Expression level changes (before and after the exposure) of microRNA levels in blood samples from rats (at least three) before and after exposure to ricin, O_3_, endotoxin and CO_2_. The blood samples were taken before exposure and 10 minutes after exposure. SD represents a standard deviation value. “*” indicates significant difference at *p-value*<0.05.

| microRNA | Ricin | | Endotoxin | | Ozone | | Carbon dioxide | |
| --- | --- | --- | --- | --- | --- | --- | --- | --- |
|  | Expression level change | SD | Expression level change | SD | Expression level change | SD | Expression level change | SD |
| miR-122 | 1.03 | 2.73 | -0.70 | 0.19 | 5.22 | 8.50 | -0.27 | 0.41 |
| miR-125b | 1.04 | 2.74 | 1.85 | 4.21 | 0.87 | 1.66 | -0.26 | 0.17 |
| miR-146a | 0.37 | 1.14 | -0.37 | 0.59 | 0.52* | 0.26 | 0.48 | 0.71 |
| miR-155 | 2.59 | 3.47 | 4.24 | 5.01 | -0.47* | 0.07 | -0.26 | 0.04 |
| miR-20b | -0.53 | 0.47 | -0.81 | 0.16 | 0.48 | 0.69 | 0.65 | 0.64 |
| miR-210 | 0.24 | 1.66 | -0.68 | 0.33 | 2.07 | 2.69 | 0.12 | 0.17 |
| miR-21 | 0.46 | 1.77 | -0.36 | 0.55 | -0.11 | 0.59 | 0.65 | 0.46 |
| miR-33 | -0.43* | 0.17 | -0.60 | 0.35 | 1.64 | 2.17 | 0.23 | 0.30 |

**Video S1.** Video clip of rat with the exposure to ozone for 10 minutes.
